# From omics to enhanced fungal virulence: Overexpression of a putative secreted protein improves *Beauveria bassiana* biocontrol potential against the insect pests *Piezodorus guildinii* and *Tenebrio molitor*

**DOI:** 10.1101/2025.04.02.646755

**Authors:** Héctor Oberti, Lucia Sessa, Emmeline van Roosmalen, Charissa de Bekker, Michael F. Seidl, Andrea Sanchez-Vallet, Eduardo Abreo

**Author notes:** Corresponding authors: Eduardo Abreo; Héctor Oberti.

## Abstract

The entomopathogenic fungus *Beauveria bassiana* is widely used as a biocontrol agent, but its efficacy varies depending on the target insect species*. Piezodorus guildinii*, a major soybean pest, exhibits low susceptibility to *B. bassiana*. Thus, biocontrol of this pest requires improving virulence of *B. bassiana*. Here, we used genomic and transcriptomic analyses to identify novel genes associated with the enhanced virulence of *B. bassiana* strain ILB308 when exposed to the insect epicuticular hydrocarbon n-pentadecane. Comparative analysis with the lower-virulence strain ILB205 revealed differentially expressed genes (DEGs) linked to genetic information process pathways and immune evasion. Among these DEGs, we identified *BbCBM9_1*, a secreted carbohydrate-binding protein, as a candidate virulence factor. Overexpression of *BbCBM9_1* in ILB205 led to enhanced virulence against *P. guildinii* and *Tenebrio molitor*, as well as improved conidial germination, tolerance to oxidative and cell wall stress, and increased growth in nutrient-rich and glycerol-supplemented media. This demonstrates that the enhanced virulence observed in ILB308 grown on n-pentadecane can be replicated by overexpressing an associated gene in a low-virulence strain. Our study, highlights the potential of integrating transcriptomics and targeted genetic modifications to optimize fungal biocontrol agents for improved pest management.

## Introduction

*Beauveria bassiana* is a well-known entomopathogenic fungus widely used as a biocontrol agent against various insect pests, including mosquitoes (*Anopheles* spp.), moths (*Spodoptera* spp.), and beetles (*Coleoptera* spp.) (Skinner *et al*., 2014). However, some insect species like the red-banded stink bug *Piezodorus guildinii* are less prone to be infected by *B*. *bassiana* (Zerbino *et al*., 2015)*. Piezodorus guildinii* is one of the primary insect pests affecting soybean crops in Argentina, Brazil, Uruguay, and USA (Sosa-Gómez *et al*., 2020). It is increasingly tolerant to chemical insecticides in the field, reducing the range of effective management strategies (Castiglione *et al*., 2008; Parys and Portilla, 2020). Similar resistance to conventional control strategies has been also observed in other important agricultural pests such as the diamondback moth *Plutella xylostella* (APRD, 2021) and the cattle tick *Rhipicephalus microplus* (Obaid *et al*., 2022). This increases the urgency to improve biocontrol agents such as *B. bassiana* to establish reliable and sustainable pest management strategies based on their enhanced efficacy against insect populations.

At the start of the infection cycle, *B. bassiana* conidia adhere to the insect’s epicuticle (outermost cuticular layer), germinate, and penetrate both the epicuticle and cuticle to breach into the insect’s hemocoel (Ortiz-Urquiza and Keyhani, 2013; Pedrini *et al*., 2013; Valero-Jiménez *et al*., 2016). To degrade the cuticle, *B. bassiana* secretes proteins such as chitinases (Fang *et al*., 2005), Pr1 family subtilisin-like proteases (Gao *et al*., 2020), and endo-β-1,3-glucanases (Wang *et al*., 2023). Additionally, it secretes effector proteins to evade insect immune responses such as LysM domain-containing proteins (Cen *et al*., 2017) as well as small, secreted proteins (Mou *et al*., 2021). Most of these proteins contain carbohydrate-binding modules (CBMs), like CBM1 (cellulose-binding module), CBM18 (chitin-binding module), or CBM50 (LysM module) (Cen *et al*., 2017; Bhagwat *et al*., 2021) that are associated with increased substrate binding and enhanced activity (Junges *et al*., 2014; Bhagwat *et al*., 2021). This suggests that secreted CBM-containing proteins could be potential targets for genetic engineering to improve biocontrol efficacy.

The composition of the cuticle, mainly consisting of chitin and proteins with a layer of lipids and hydrocarbons, varies depending on the insect species (Pedrini *et al*., 2013). In *P. guildinii*, n-pentadecane (HC15) is a major epicuticular component (Sessa *et al*., 2021). Exposure to this compound has been found to enhance the virulence of *B. bassiana* strains, such as ILB308, while other strains, like ILB205, do not respond (Sessa *et al*., 2022). Based on this observation, we hypothesized that identifying genes related to this enhancement could provide valuable insights for genetic improvement of low-virulence strains that have other traits of interest, including increased conidiation capacity or high tolerance to abiotic stress. To explore this hypothesis, we performed a comparative transcriptomic analysis of *B. bassiana* strains ILB205 and ILB308 to identify candidate genes responsible for the HC15-mediated virulence enhancement. We then proceeded to functionally validate a promising candidate gene encoding a secreted CBM-containing protein. Jointly, these findings underscore the potential of integrating transcriptomics with targeted genetic modifications to develop more effective fungal biopesticides for pest management.

## Experimental Procedures

### Biological materials

*B. bassiana* strains ILB205 and ILB308 were obtained from the INIA Las Brujas fungal collection, Uruguay (WDCM 1291), and grown on potato dextrose agar (PDA) (Sigma, St. Louis, USA) plates for 7 days at 25 °C.

*Escherichia coli* One Shot TOP10 (ThermoFisher Scientific, Waltham, Massachusetts, USA) was grown in Luria Broth (LB) (Sigma) for 1 day at 37 °C and employed for DNA manipulations and transformations.

*Agrobacterium tumefaciens* AGL-1 was used for fungal transformations and grown in LB supplemented with carbenicillin (100ug/ml) (Sigma) and rifampicin (50ug/ml) (Sigma) for 2 days at 28 °C.

Individuals of *P. guildinii* were obtained from an indoor mass rearing and maintained at 24 ± 1 °C, 80 ± 10% relative humidity, and 14 h of photo phase on fresh soybeans, green beans, and water. Adults were sexed and placed in plastic cages prior to each bioassay.

Adult individuals of yellow mealworm beetle *Tenebrio molitor* of 1-2 weeks old were obtained from “La Biofabrica” (Montevideo, Uruguay) and maintained no longer than 1 week at 25 °C in the dark in plastic boxes.

*Piezodorus guildinii* exuviae of 5th instar nymphs were obtained from an indoor mass rearing. The collected exuviae were first autoclaved and then dried to constant weight in an oven at 40 °C (Sessa *et al*., 2024). These exuviae were used as a proxy for epicuticle of the insect, which is the outer layer of the cuticle and comprises a heterogeneous mixture of lipids that include abundant levels of long-chain alkanes, alkenes, wax esters and fatty acids (Pedrini *et al*., 2010).

### Fungal growth conditions and RNA-seq data acquisition

*Beauveria bassiana* strains ILB308 and ILB205 were grown in different axenic culture conditions for RNA extraction (**Supplementary Figure 1, Supplementary Table 1**): (i) Minimal media (MM) (0.4 g/L KH2PO4, 1.4 g/L Na2HPO4, 0.6 g/L MgSO4⋅7H2O, 1.0 g/L KCl, 0.7 g/L NH4NO3⋅7H2O and 15 g/L agarose)-this condition was performed only for ILB205, ILB308 data was downloaded from NCBI, see above-; (ii) MM supplemented with *P. guildinii* epicuticle as the only carbon source (MM+EC); or (iii) conidia previously grown on MM supplemented with 10% v/v HC15 (Acros Organics, ThermoFisher Scientific, Fair Lawn, NJ, USA), harvested, and then cultured on MM+EC plates (HC15_MM+EC). In all EC conditions 40 mg of *P. guildinii* epicuticle was added at the top layer of the culture media so it could be in direct contact with *B. bassiana* strains growing on it. For each condition, three biological replicates were incubated at 25 °C for 4 days. Fungal mycelia were collected from each sample (replicate) and RNA extraction was conducted. RNA-seq data of strain ILB308 in Minimal Medium (MM) as well as infection at 4 days post inoculation (dpi) of *P. guildinii* with and without pre-growth in MM supplemented with HC15 were obtained from NCBI Bioproject PRJNA1117794 (Oberti *et al*., 2025) (**Supplementary Table 1**).

For RNA extraction, each fungal sample was ground with an RNAse-free mortar and pestle in liquid nitrogen. Total RNA was extracted using the RNAeasy Plant isolation kit (Qiagen, Germany). Polyadenylated mRNA was strained from the total RNA and cDNA libraries were prepared using the Truseq RNA Library Prep mRNA kit (Illumina, San Diego, USA) following the manufacturer’s instructions. Constructed libraries were sequenced at 150×2 bp PE with an Illumina Novaseq6000 (Illumina) at Macrogen Inc, Korea.

### RNA-seq expression analysis

Raw RNA-seq data was filtered using Trimmomatic v0.36 (Bolger *et al*., 2014) to trim Illumina adapters and bases with a quality score value less than 25 from both sides of read ends; final minimum size allowed after trimming was 75 bp. For all expression analyses, the *B. bassiana* strain ILB308 genome assembly (NCBI Acc JBEBMD000000000) was used as a reference. Structural and functional annotations of this genome were previously obtained in Oberti *et al*., 2025. Filtered reads were aligned to the reference genome using STAR v2.7 (Dobin *et al*., 2013) and subsequently summarized with RSEM v1.2.8 (Li and Dewey, 2014) to determine read counts per gene. All statistical analyses of the expression data were performed using EdgeR v3.28 (Robinson *et al*., 2009). Differentially expressed genes (DEGs) were identified using the exact test with the following conditions: CPM>1 in at least two repetitions, log2FoldChange (FC) of |1|, and FDR p< 0.05. Functional annotation of all genes of GO terms and KEGG pathways was used for enrichment analysis of DEGs using the Fisher Exact test and FDR p< 0.05 against all annotated genes.

### Candidate gene selection and *in silico* analysis

Given the role of secreted, CBM-containing proteins in the infection process of entomopathogenic fungi, and the observed enhancement in virulence of ILB308 following HC15 treatment, we established the following criteria for selecting the candidate genes: (i) encoded a secreted protein, as described in Oberti *et al*., 2025; (ii) contained at least one CBM module, (iii) was up-regulated after HC15 treatment in ILB308 during growth on *P. guildinii* epicuticle, and (iv) was up-regulated during infection of *P. guildinii* at 4 dpi.

To investigate candidate gene conservation across different genera of fungi, we performed an orthology analysis using Orthofinder2 (Emms and Kelly, 2019). A set of protein-coding genes of 18 publicly available structural annotations of fungi comprising entomopathogens, plant pathogens, endophytes, symbionts, and mycoparasites were obtained from NCBI (**Supplementary Table 2**). Orthologous gene pairs were identified based on amino acid sequence similarity and applying a coverage cutoff of 50% of the total length of the shorter gene sequence. Sequence similarity was established with BLASTp v2.16 (Altschul et al., 1990) and a threshold E-value 1e^-5^.

Sequence analysis of up and downstream regions of candidate gene was done by mapping ILB205 whole genome sequences (obtained from NCBI Project PRJNA1117794) to the ILB308 genome using BWA 0.7.17 (Li, 2013). RNA and DNA sequence data mapping was visualized using IGV genome browser (Thorvaldsdóttir *et al*., 2013).

### Generation of overexpression mutants

First, the general “over-expression vector” pER023 was constructed to facilitate easy swapping of genes of interest for functional analysis in *B. bassiana*. As a backbone we used vector LIIIβ F A-B, BB53 (Binder *et al*., 2014), which holds the needed genetic information for vector cloning into *E. coli* followed by *Agrobacterium*-mediated transformation. This backbone contains *E. coli* and *Agrobacterium* origins of replication, a kanamycin resistance cassette, left and right T-DNA borders and a resolvase. Using NEBuilder HiFi DNA Assembly (NEB, US) the EcoRV restriction site in the backbone was removed and a bialaphos resistance (bar) cassette (Pall and Brunelli, 1993) was incorporated in between the T-DNA borders for phosphinothricin-based transformant selection. An expression cassette under the control of the constitutive *TrpC* promoter and *TrpC* terminator originating from *Aspergillus nidulans* was also placed in between the T-DNA borders. An EcoRV restriction site (GATATC) was incorporated between the promoter and terminator for easy incorporation of genes of interest.

The open reading frame (ORF) of the candidate gene *BbCBM9_1* was amplified from ILB308 gDNA using Q5 Polymerase (NEB, US) with specific primers (**Supplementary Table 3)**. Binary vector pER023 was linearized using EcoRV (NEB, US) and the ORF of *BbCBM9_1* was incorporated using NEBuilder HiFi DNA Assembly (NEB, US). The resulting vector was purified using the GeneJET Plasmid Miniprep Kit (ThermoScientific, USA) and sequenced for verification by Plasmidsaurus Inc, USA (**Supplementary Figure 2**). The verified final vector was transformed into *A. tumefaciens* AGL-1 using the triparental mating method (Raman and Mysore, 2023).

*S*train ILB205 was selected as wild type strain for the generation of overexpression mutants due to its low-virulence (compared to ILB308) on *P. guildinii*. Transformation of *B. bassiana* was carried out as previously described in Oberti *et al*., 2025 with slight modifications. *Agrobacterium tumefaciens* strain AGL-1 was grown at 28°C at 200 rpm for 18 h in liquid LB medium supplemented with kanamycin (50 ug/ml) (Sigma), rifampicin (50ug/ml) (Sigma), and carbenicillin (100ug/ml) (Sigma). The culture was diluted to an optical density 660 nm (OD660) of 0.15 in induction medium (IM) - 10 mM K2HPO4,10mM KH2PO4, 2.5 mM NaCl, 2 mM MgSO4, 0.7 mM CaCl2, 9 mM FeSO4, 4mM NH4SO4, 10 mM glucose, 40 mM 2-[N-morpholino] ethanesulfonic acid, pH 5.3, 0.5% glycerol (w/v), 200 mM acetosyringone. Cells were grown under these conditions until an OD660 of 0.6–0.8 was reached before mixing them with an equal volume of a fresh conidial suspension of ILB205 (1×10^6^ conidia/ml). Conidia were harvested from PDA plates growth for 7 days at 25 °C. This mix was plated on solid IM (IM with 15 g/l agar). After co-cultivation at 28°C for 48 h, a layer of top-agar of Czapek-Dox media (30 g/L sucrose, 2 g/L NaNO3, 1 g/L K2HPO4, 500 mg/L KCl, 500 mg/L MgSO4, 10 mg/L FeSO4, and 7.5 g/L agar) supplemented with phosphinotricin (300ug/ml) (Sigma) as the selection agent for fungal transformants and cefotaxime (600 ug/ml) (Sigma) to inhibit growth of AGL-1 *A. tumefaciens*. Two independent colonies from different plates were selected, and monosporic cultures of each were obtained and grown again in selection medium. To determine the stability of the two monosporic transformant lines, they were successively cultured on PDA for five generations, after which transformants were transferred to Czapek-Dox supplemented with the selection agent again.

To evaluate the relative expression of *BbCBM9_1* in these overexpression mutants compared to the wild-type strain, primers were designed for this gene, while the *glyceraldehyde-3-phosphate dehydrogenase* (*GAPDH*) gene from *B. bassiana* (Mantilla *et al*., 2012) was used as a reference gene (**Supplementary Table 3**). To assess primer amplification efficiency, standard curves were constructed by plotting the log of cDNA dilution values versus CT values. Serial 5-fold dilutions of cDNA samples were used as templates for each qRT-PCR reaction. Each reaction efficiency was calculated as E = 10 (−1/slope), using the slope of each curve. For expression quantification RNA extractions were performed from PDA 4-day old grown cultures at 25 °C. Total RNAs were extracted from the resultant cultures with the RNeasy plant mini kit (Qiagen). RNA was quantified using Nanodrop (Thermo Fisher) and integrity and purity were evaluated on an agarose gel 1% electrophoresis. All samples were treated with Turbo DNase (Invitrogen, Thermo Fisher Scientific) and EDTA. Purified RNA was transcribed into cDNA using iScript cDNA Synthesis Kit (Bio-Rad). qRT-PCR analyses were performed using iQ SYBR Green Supermix (Bio-Rad). PCRs were performed using Quant Studio 3 (Applied Biosystems, Thermo Fisher), using the following conditions: denaturation at 95 °C for 3 min, followed by 40 cycles of 95 ◦C for 15 s and 60 ◦C for 30 s. Data was analyzed using the Quant Studio design and analysis software version 1.4.3. The qRT-PCR analyses were performed in triplicate for each of the three independent biological replicates using the wild-type strain samples as a control. Relative expression levels of candidate gene were calculated using the 2−ΔΔCt method (Livak and Schmittgen, 2001).

### Virulence Assays

Virulence assays were conducted using *P. guildinii* and *T. molitor*, a model insect that is widely used in entomopathogenic fungal bioassays (Bharadwaj and Stafford, 2011). Conidia of the wild type strain and overexpression mutants were harvested as detailed above from a 7 days old PDA culture. Insects were inoculated by immersion for 10 s in each spore suspension (1×10^7^ conidia/mL), placed individually on a 55 mm Petri dish containing sterile paper, and maintained under controlled conditions (28°C, 70% relative humidity). Mortality was corrected for control effects by using Abbot’s formula (Abbott, 1925). Corrected cumulative survival curves were constructed and plotted. Lethal time LT50 (time taken to kill 50% of insects) values were determined using R custom scripts and results were compared using a one-way ANOVA for determining the differences between treatments and means were compared by Tukey test (95%) using GraphPad Prism version 9 for Windows (GraphPad Software Inc., San Diego, CA, USA; www.graphpad.com).

### Phenotypic evaluations of transformed strains

Wild type and overexpression mutant strains were evaluated based on growth in different carbon sources, stress tolerance, and conidial viability. Conidia suspensions used in all experiments were harvested from 1-week-old PDA plates.

For growth assays in different carbon sources, 2 μl of a 1×10 conidia/ml suspension were centrally spotted onto PDA and Czapek-Dox media, where sucrose (3% w/v) was replaced with 3% of glucose, fructose, maltose, trehalose, glycerol, or mannitol. Colony diameters were measured after 7 days of growth at 25°C.

For stress tolerance assays, the same conidial suspensions were spotted on Czapek-Dox media supplemented with stress-inducing agents: (i) osmotic stress (0.8 M NaCl, 1 M sorbitol), (ii) oxidative stress (2 mM H₂O₂), and (iii) cell wall stress (4 μg/ml Congo Red, 5 μl/ml Calcofluor White). Colony diameters were recorded after 7 days of growth at 25°C.

Conidial viability was assessed by spreading 150 μl of a 1×10 conidia/ml suspension onto 2% water agar plates, incubating for 18 hours at 25°C in darkness and counting germinated spores of a total of 300 spores per plate.

All experiments were performed in triplicate with three biological replicates. One-way ANOVA was used for determining the differences between treatments and means were compared by Tukey test (95%) using GraphPad Prism version 9 for Windows (GraphPad Software Inc., San Diego, CA, USA; www.graphpad.com).

## Results

### *B. bassiana* strains ILB205 and ILB308 respond differentially to epicuticle interaction

Despite their close phylogenetic relationship (Héctor Oberti *et al*., 2025), *B. bassiana* strains ILB205 and ILB308 show differences in virulence towards *P. guildinii* after exposure to HC15: while the virulence of ILB205 is not affected by HC15, virulence of ILB308 increases significantly (Sessa *et al*., 2022). We designed a transcriptomic experiment to identify genes that are specifically affected by exposure to HC15 and their subsequent interaction with the epicuticle of *P. guildinii* in ILB308. To this end, we sequenced the transcriptomes of these two strains in three different growth conditions; MM, MM+EC and HC15_MM+EC (**Supplementary Figure 1**). We obtained a total of 52.1 Gigabases (Gb) of sequencing data (on average 2.9 Gb per sample) (**Supplementary Table 1**). We mapped all RNAseq data to the ILB308 reference genome assembly (H. Oberti *et al*., 2025) (**Supplementary Table 1**) and subsequently performed differential gene expression analysis. Pairwise comparisons were made between strains ILB205 and ILB308 under the same growth conditions. We compared i) ILB308 MM with ILB205 MM, ii) ILB308 MM+EC with ILB205 MM+EC, and iii) ILB308 HC15_MM+EC with ILB205 HC15_MM+EC (**Figure 1A, Supplementary Figure 1**). Based on these three pairwise comparisons, we identified a total of 3,658 DEGs, which represent 40% of all predicted protein-coding genes in the B. *bassiana* strain ILB308. The condition with the highest number of DEGs was MM+EC, with 2,754 genes (**Figure 1A**), highlighting that ILB308 and ILB205 exhibit significantly different responses to growth on *P. guildinii* epicuticle.

**Figure 1.**
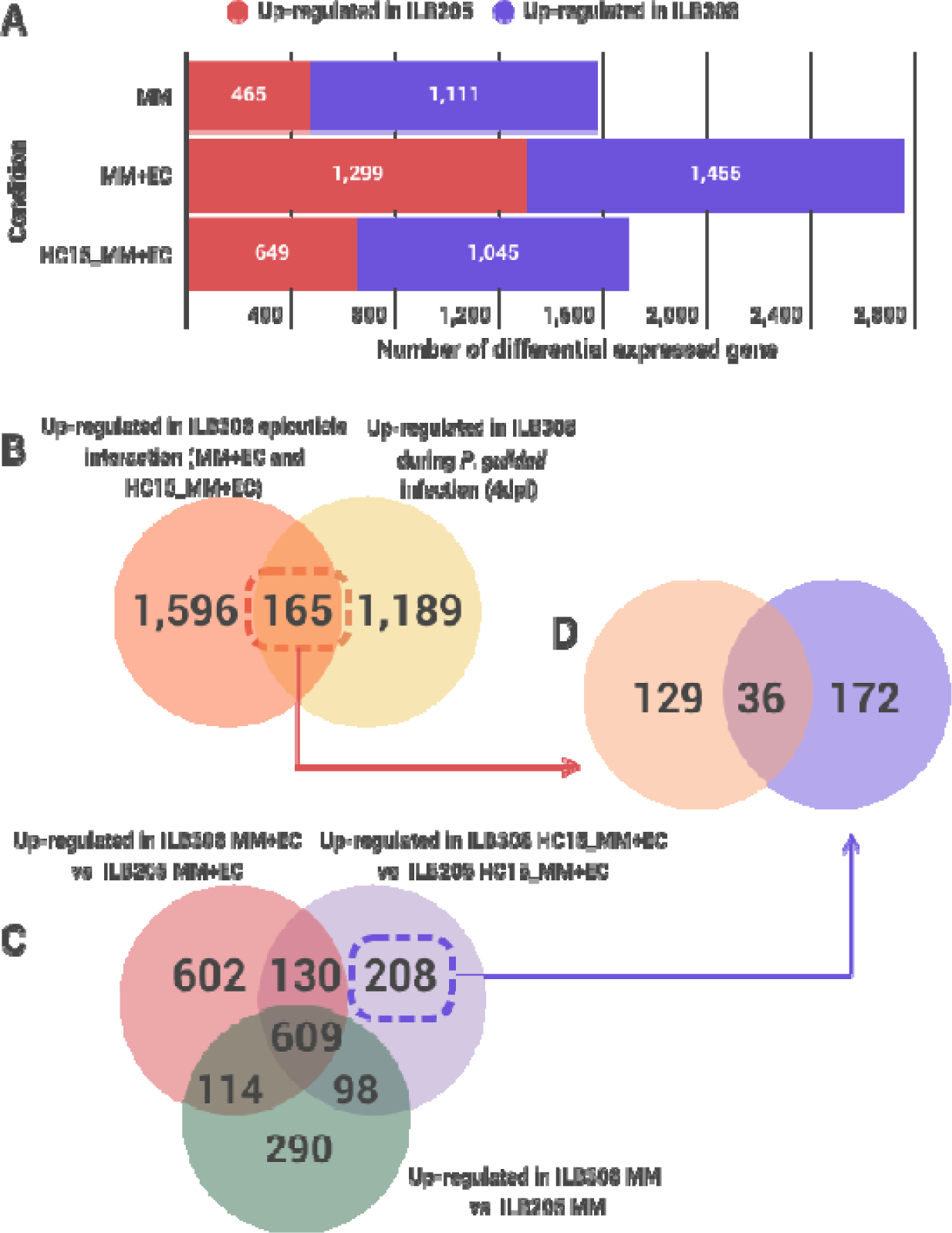
Transcriptome analyses reveal different responses of ILB205 and ILB308 to *P. guildinii* epicuticle. A) Number of differentially expressed genes in *B. bassiana* strain ILB308 compared to ILB205. Differentially expressed genes were identified at a logFC >|1| and a false discovery rate (FDR) < 0.05 in pairwise comparisons. Venn diagrams show ILB308 (B) up-regulated genes during epicuticle interaction compared to ILB205 and during infection after 4 days post inoculation (versus MM; obtained from Oberti et al 2025); (C) up-regulated genes compared to ILB205 in all analyzed conditions; and (D) combination of genes that were up-regulated during epicuticle interaction, infection of *P. guildinii* and exclusively up-regulated after HC15 growth. The red rectangle highlights the number of shared up-regulated genes between epicuticle interaction and infection at 4dpi. The blue rectangle highlights exclusively up-regulated genes in HC15_MM+EC conditions. MM: Minimal medium; MM+EC: MM supplemented with *P. guildinii* epicuticle; HC15_MM+EC: conidia pre grown in MM supplemented with n-pentadecane (HC15), harvested and then grown in MM+EC.

When comparing differences in expression between ILB308 and ILB205 during interaction with *P. guildinii* epicuticle, we found that 1,455 and 1,045 genes were up-regulated in ILB308 in MM+EC and HC15_MM+EC, respectively, (**Figure 1A)** accounting for a total of 1,761 genes. Among these genes, we found chitinases, chitin synthase, 1,3-beta-glucanosyltransferas, beta-glucanases and cytochrome P450 monooxygenases (CYP450) (**Supplementary Table 4)**, suggesting that both strains have a clear induction of different genes after epicuticle interaction, that could contribute to differences in cuticle degradation, nutrition acquisition, or stress tolerance, thereby affecting virulence.

To identify functional enrichment in each set of up-regulated genes in ILB308, we performed GO and KEGG pathway enrichment analyses. The GO terms ‘RNA-directed DNA polymerase activity’, ‘RNA-templated DNA biosynthetic procesś, and ‘DNA polymerase activity’ were enriched (**Supplementary Table 5**). Similarly, the KEGG pathways ‘Homologous recombination’, ‘Non-homologous end-joining’, and ‘Cysteine and methionine metabolism’ (**Supplementary Table 5**) were significantly enriched. These analyses show that differences between ILB308 and ILB205 related to EC interaction might involve genes with putative roles in RNA biosynthesis and genetic information processing that could be associated with unique features of ILB308.

Up-regulation of genes related to host immune evasion and hydrocarbon assimilation are shared during epicuticle interaction and *P. guildinii* infection.

We have previously reported 1,354 up-regulated genes in ILB308 during infection of *P. guildinii* at 4 dpi compared to MM (H. Oberti *et al*., 2025). We observed that 165 of these genes were also up-regulated in ILB308 during the interaction with *P. guildinii* epicuticle (MM+EC and HC15_MM+EC) (**Figure 1C, Supplementary Table 4**). Among these genes, we found two homologues to previously functionally verified genes involved in *B. bassiana* virulence: *BbILB308_006964, BbILB308_08180.* These genes encode for the secreted LysM domain containing protein *Bly8 (B. bassiana* ARSEF 2860 gene *BBA_09350)* (Cen *et al*., 2017), a small, secreted cysteine-free protein (*cfp*) *(B. bassiana* ARSEF 2860 gene *BBA_ 02121)* (Feng *et al*., 2015) respectively. We also found three cytochrome CYP450 encoding genes, probably related to assimilation of n-alkanes like HC15 as carbon sources (Pedrini *et al*., 2013) (**Supplementary Table 4**). These observations suggest that differences in virulence between ILB308 and ILB205 could be related to different expression patterns of genes related to hydrocarbon assimilation and host defense evasion.

### Secreted proteins are up-regulated after pre-growth in n-pentadecane

Since only ILB308 shows enhanced virulence when exposed to HC15, we hypothesized that genes exclusively up-regulated in this condition compared to ILB205 might contain key genes responsible for this unique response. Our analysis revealed 208 genes up-regulated solely in HC15_MM+EC in ILB308 (**Figure 1C, Supplementary Table 4**), with 36 of these genes also up-regulated during infection of *P. guildinii* (**Figure 1D, Supplementary Table 4**). Within these 36 genes, a clock-controlled protein exhibited the highest expression levels in HC15_MM+EC (**Figure 2**). Notably, none of these 36 genes have been functionally characterized in existing databases, which underline their novel roles in HC15-potentiated pathogenic responses.

**Figure 2.**
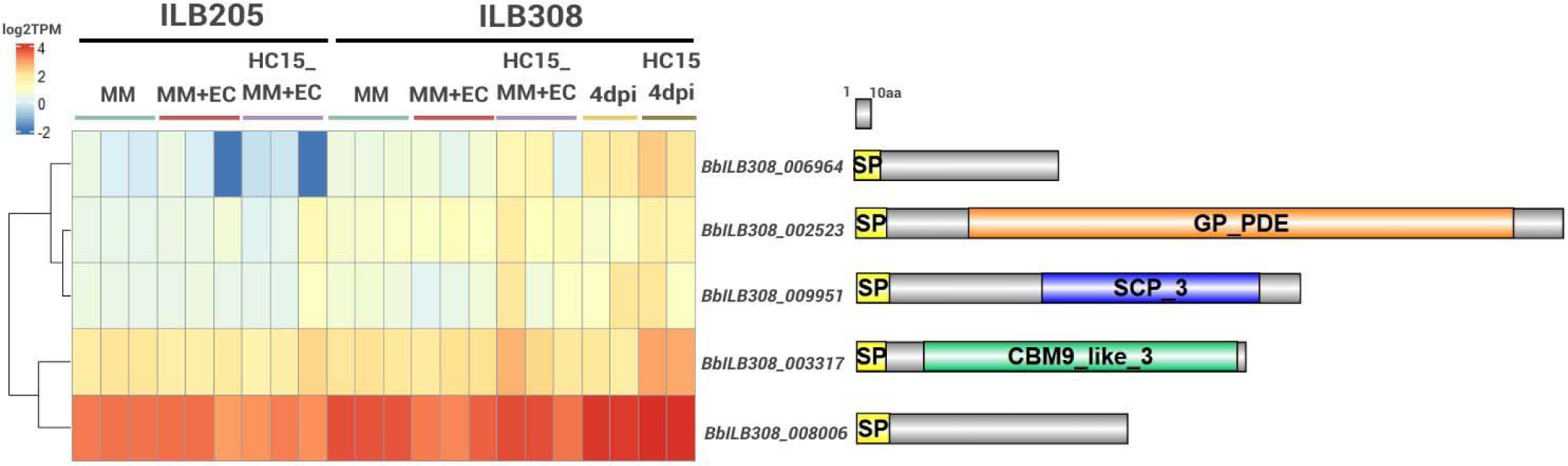
Heatmap and protein domains of five putative secreted genes specifically overexpressed in ILB308 during interaction with *P. guildinii* epicuticle after pre growth in HC15. Each row represents a single gene, and each column corresponds to a sample. Expression values are depicted by a color scale ranging from dark blue (high expression) to red (low expression). Genes were clustered based on expression similarity using hclust complete clustering algorithm, with results visualized through a dendrogram. Protein illustration and domains were drawn next to each gene. Conditions include Minimal Medium (MM), Minimal Medium supplemented with *P. guildinii* epicuticle (MM+EC), pre-growth in HC15 followed by growth in MM+EC (HC15_MM+EC), infection of *P. guildinii* at 4 days post-inoculation (4dpi), and infection of *P. guildinii* at 4 dpi after fungal growth on HC15 (HC15 4dpi). SP: signal peptide; CBM9_like_3: Carbohydrate binding module 9 like 3; SCP_3: Bacterial SCP orthologue; GP_PDE: Glycerophosphodiester phosphodiesterase domain

Given the relevance of secreted proteins in fungal pathogens (eg: Lu and Edwards, 2016; Cen *et al*., 2017; Mou *et al*., 2021, 2022) we cross-referenced these 36 candidates with those previously predicted as putatively secreted in ILB308 (Oberti *et al*., 2025). This analysis identified five secreted proteins (*BbILB308_002523*, *BbILB308_003317, BbILB308_006964, BbILB308_008006,* and *BbILB308_009951*), of which only one contained a CBM domain (*BbILB308_003317) (***Figure 2**). While gene *BbILB308_003317* was similarly expressed in all conditions, it was up-regulated in ILB308 after HC15 pre-growth during epicuticle interaction and infection (**Figure 2**). This up-regulation following HC15 exposure indicates that *BbILB308_003317* reacts to this compound and could, therefore, play a role in the enhanced virulence.

### Sequence analysis of the conserved *BbCBM9_1* protein

*BbILB308_003317* is 807 bp in length with no predicted introns and has a coding sequence of 607 bp, encoding a protein with 228 amino acids (molecular mass 25.163 kDa). The predicted *B. bassiana* protein contains a signal peptide of 15 aa based on SignalP (Almagro Armenteros *et al*., 2019), which was corroborated by WoLFPSORT (Horton *et al*., 2007). This protein has a DOMON-like type 9 carbohydrate binding module (CBM9_like_3) in position 52-227aa (**Figure 2)**. It is the only protein with a CBM9 domain in the entire predicted proteome of ILB308, according to InterproScan (Jones *et al*., 2014), and thus we named it *BbCBM9_1*. The ILB308 *BbCBM9_1* gene is homologous to the characterized secreted protein XP_008599210.1 (99.6% amino acid similarity) of *B. bassiana* ARSEF2860 (Santi *et al*., 2018) and to CFP28 (60.5% amino acid similarity), a DOMON-like type 9 carbohydrate-binding module domain-containing protein of the fungal pathogen *Coccidioides posadasii* (Lunetta and Pappagianis, 2014). The sequence similarity to these two secreted proteins further strengthened the prediction that *BbCBM9_1* is indeed a secreted protein.

We also checked for conservation of *BbCBM9_1* among eight *B. bassiana* strains and across 18 well annotated fungal species with different lifestyles. We found that this gene was conserved in all *B. bassiana* strains as a single-copy gene. Orthologous proteins were identified in 15 of the 18 fungal species analyzed (**Supplementary Table 6**). Orthologs of *BbCBM9_1* were absent in *Claviceps purpurea*, a phytopathogen, and two entomopathogens, *Hirsutella minnesotensis* and *Ophiocordyceps sinensis*. Additionally, while *BbCBM9_1* generally occured as a single copy in 13 of the analyzed species, we observed two exceptions: *Tolypocladium ophioglossoides*, a mycoparasitic fungus, and *Pochonia chlamydosporia*, a nematophagous fungus, which both had two copies of the gene (**Supplementary Table 6)**. Sequence analyses of the *BbCBM9_1* orthologues showed that the CB9M-like domain as well as the signal peptide are highly conserved. This degree of conservation across different fungal species with different lifestyles suggests that this gene was likely present in the common ancestor of these species and might have a conserved function.

We observed that *BbCBM9_1* differs between strains ILB205 and ILB308 by a single nucleotide substitution (**Supplementary Figure 3**), resulting in an amino acid change at position 142 (Val to Ile) that occurs within the CBM9_like_3 domain. To explain differences in gene expression, we also searched for nucleotide differences in the presumed promoter region of the gene up to 2kb upstream. We observed that 99.5% of the nucleotides in this promoter region were identical, with differences occurring only in eleven positions (**Supplementary Figure 3**). Moreover, no transposable elements (TE) were detected 10 kb up-or downstream of this gene in both strains, making it unlikely that the presence of these elements impact the expression of *BbCBM9_1*. These observations collectively indicate that differences in gene expression between strains may instead be dueto regulatory interactions involving distant enhancer elements, or other regulatory genes such as transcription factors specific to each strain.

### Overexpression of *BbCBM9_1* improves virulence towards *T. molitor* and *P. guildinii*

We hypothesized that *BbCBM9_1* plays a role in the enhancement of virulence in ILB308 after growth in HC15. To test this hypothesis, we overexpressed the gene in the less virulent strain ILB205, which does not respond to HC15 (Sessa *et al*., 2022). Two independent ectopic insertion mutants were generated, and overexpression of *BbCBM9_1* was measured by RT-qPCR. Expression analysis showed that the two mutants overexpress the gene 2.5 and 3 times compared with wild-type expression levels in PDA (**Supplementary Figure 4**), which is comparable to the 2.77 FC observed between ILB308 and ILB205 in HC15_MM+EC.

To evaluate the impact of *BbCBM9_1* overexpression on fungal virulence, we conducted insect bioassays simulating natural infection through cuticle penetration (**Figure 3**). In the case of adults of *T. molitor*, both the wildtype and the overexpression mutants were able to induce high mortality levels in mutant lines 10 days post infection (dpi) (**Figure 3A**). However, the overexpression mutant lines were able to kill *T. molitor* adults significantly (p<0.01) faster (**Figure 3B**). In adults of *P. guildnii,* the overexpression mutant lines killed a higher number of insects, from 53% in wild type to approximately 80% in 10 dpi (**Figure 3D)** and did so significantly (p<0.01) faster (**Figure 3E**). These results show that *BbCBM9_1* is involved in the enhancement of virulence of *B. bassiana*.

**Figure 3.**
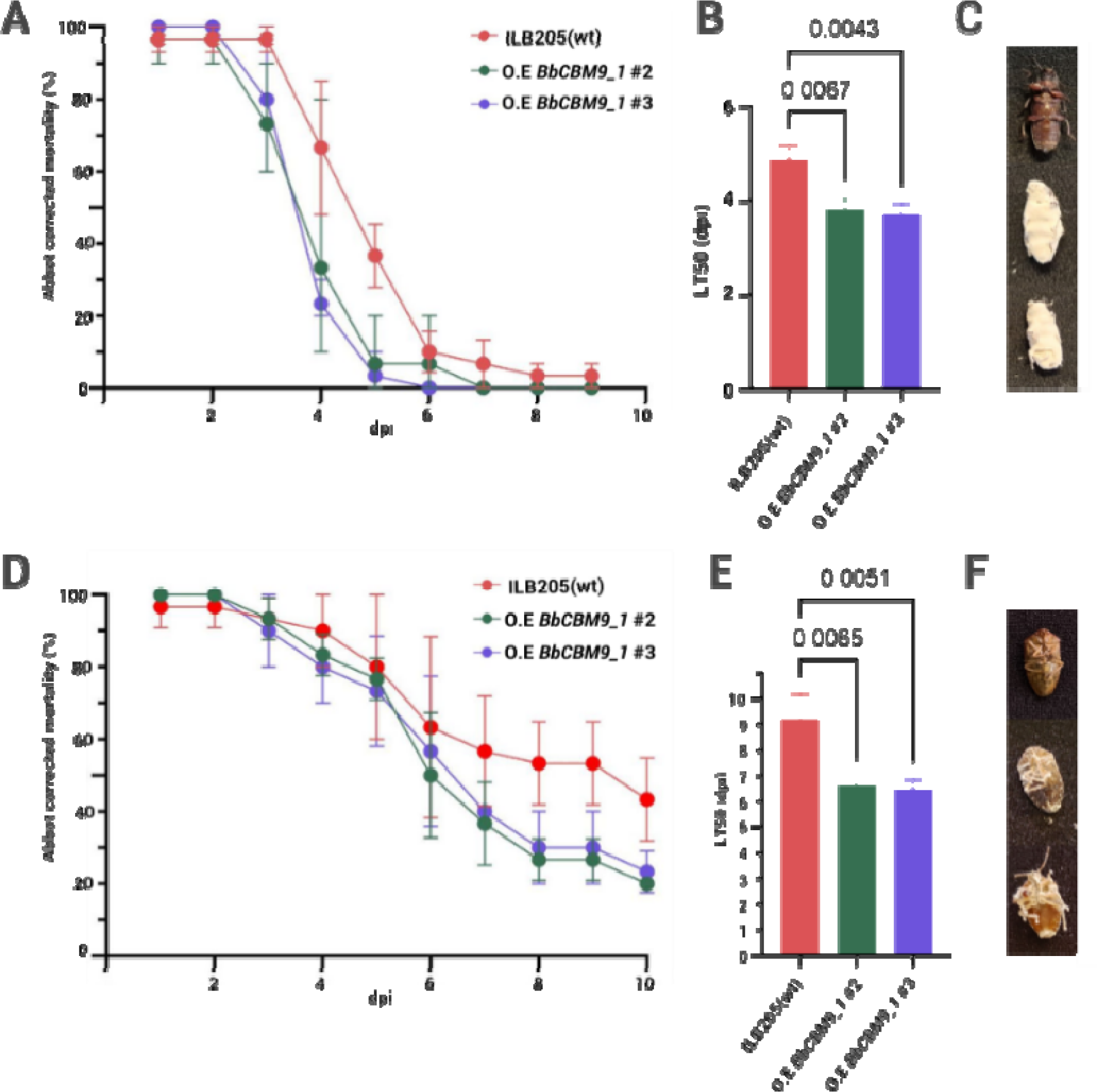
Overexpression of *BbCBM9_1* enhances virulence towards two insect species. Mortality curves and LT50 of *T. molitor* (A-B) and *P. guildinii* (D-E) adults after topical application (immersion) of a 1×10^7^ conidia/ml suspension. Illustrative photos after 20 dpi, from top to bottom of mock, ILB205, and overexpression mutant *BbCBM9_1* #2 and #3 infection of *T. molitor* (C) and *P. guildinii* (F) adults. P-values < 0.01 according to a one-way ANOVA followed by a Tukey posthoc test of results between overexpression lines versus the wild type are displayed on the plot based on triplicate experiments. Error bars: standard errors (SEs) of the means from 3 independent replicates. dpi: days post inoculation of interest strain. O.E: Overexpression mutant. LT50: time taken to kill 50% of insects. Abbot corrected mortality: mortality caused by the treatment, adjusted for natural mortality observed in the control group, using Abbott’s formula.

### *BbCBM9_1* overexpression improved conidia germination, growth in rich media and tolerance to stress

We further investigated whether the overexpression of *BbCBM9_1* had pleiotropic effects on mutant strains (**Figure 4**). For this, we evaluated radial growth differences on different carbon sources and stressors. A significant (p < 0.001) reduction in radial growth was observed exclusively in Czapek-Dox medium supplemented with trehalose, while a significant increase (p < 0.001) in growth was detected in media supplemented with glycerol and in nutrient-rich PDA medium (**Figure 4 A-B**). These findings suggest that *BbCBM9_1* may be involved in metabolic pathways related to the uptake or processing of specific carbon sources, such as glycerol and trehalose, and may also contribute to the efficient utilization of complex carbohydrates present in rich media.

**Figure 4.**
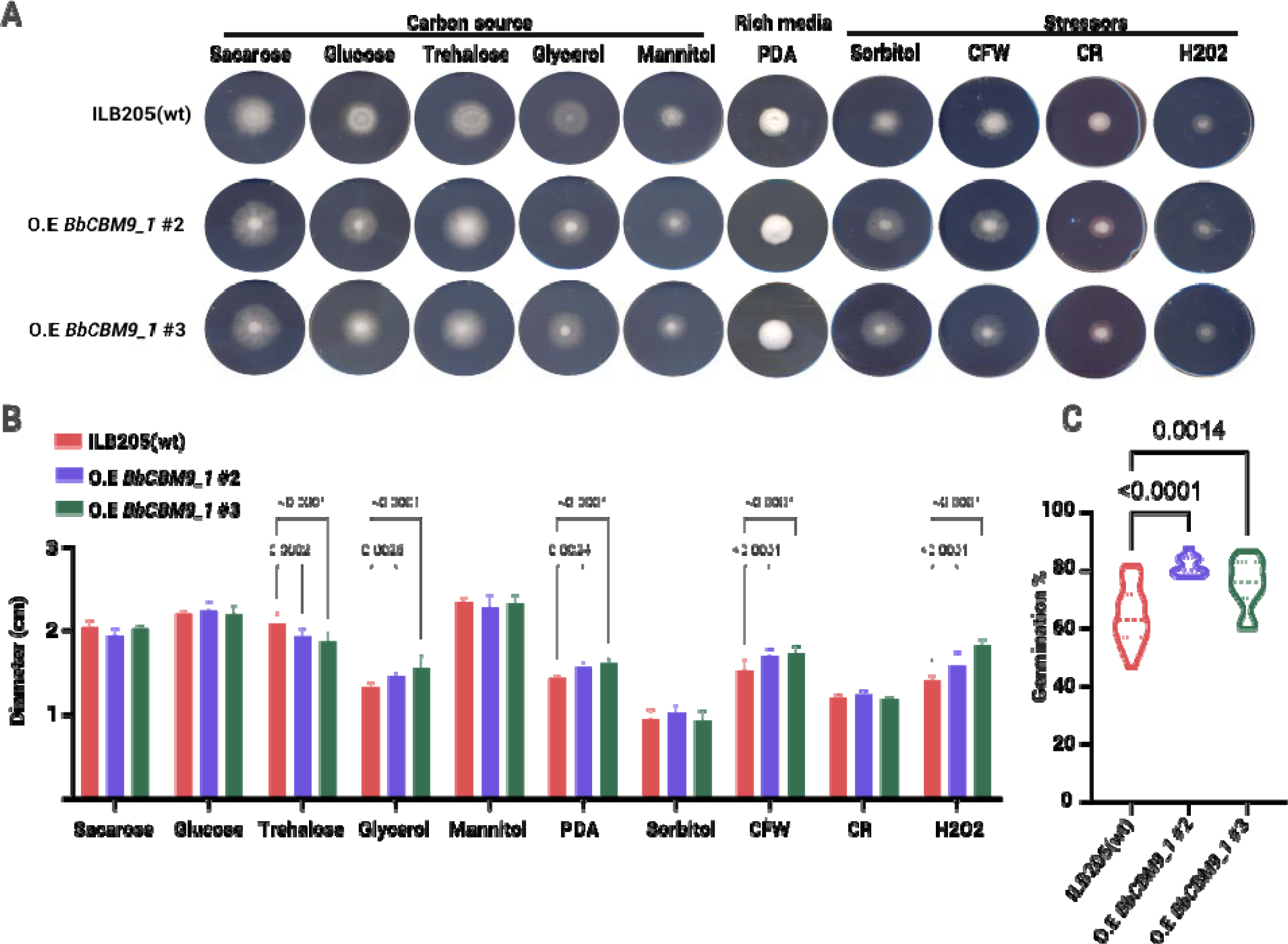
*BbCBM9_1* affects radial growth on different carbon sources, stressors and conidia germination. Images A) and diameter B) of fungal colonies incubated on Czapek-Dox media amended with different carbon sources (sucrose, glucose, trehalose, glycerol, and mannitol), rich media (PDA), osmotic agents (Sorbitol), oxidative agents (H2O2) and cell wall stressors (Calco Fluor White and Congo Red) for 7 days at 25°C. (C) Germination rate of spores harvested from PDA after 18 h in water agar. Overexpression of *BbCBM9_1* where #2 and #3 correspond to two independent overexpression mutant lines of *BbCBM9_1*. *P-values* < 0.01 according to a one-way ANOVA followed by a Tukey posthoc test of results between overexpression lines versus the wild type are display on the plot based on triplicate experiments. CFW= Calco Fluor White, CR= Congo Red.

Moreover, we tested sensitivity of overexpression mutant strains to osmotic (sorbitol), oxidative (H_2_O_2_), and cell wall stressor-Congo red and Calco fluor white (CFW)-agents on Czapek-Dox plates (**Figure 4 A-B**). No significant changes were detected for colony growth under Congo red and sorbitol. However, increased tolerance to CFW and H_2_O_2_ stressors (p < 0.01) was detected in the mutant strains compared to the wild type (**Fig. 4A-B**). These data demonstrate that the overexpression of *BbCBM9_1* improves the tolerances of *B. bassiana* to at least two types of stresses important for the adaptation to environmental changes that affect virulence.

Germination rate is a pathogenicity determinant in entomopathogens (Safavi *et al*., 2007). Consequently, we determined the effect of overexpression of *BbCBM9_1* on the germination rate of ILB205. Both mutant strains significantly increased (p<0.05) conidia germination compared to the wild type from 64% to above 75% of germinated conidia after 18 h (**Figure 4C**), suggesting that *BbCBM9_1* is involved in pathways related to germination.

## Discussion

Identifying novel genes involved in virulence enhancement is key for developing biotechnological strategies to improve *B. bassiana* strains for pest control. While certain *B. bassiana* strains exhibit relevant traits for a biocontrol agent such as high conidiation, germination, and abiotic stress tolerance, they may lack other virulence determinants against specific insect targets (Quesada-Moraga *et al*., 2024). Here, we investigate gene expression differences underlying the fungal interaction with epicuticle, virulence, and enhancement potential of two *B. bassiana* strains (ILB205 and ILB308) toward the stinkbug *P. guildinii*. Specifically, by identifying genes associated with both early cuticle interaction and later infection stages in the highly virulent strain ILB308, we sought to uncover novel potential genetic targets to be leveraged for strain improvement for targeted pest management.

First, we aimed to identify genes or pathways that differentiate the interaction of ILB205 and ILB308 with the *P. guildinii* epicuticle, focusing on ILB308 due to its higher virulence and the enhancement of its pathogenicity after pre-growth in HC15. Given that *B. bassiana* relies on cuticle degradation for host invasion, we expected ILB308 to upregulate genes linked to hydrocarbon metabolism, fatty acid processing, as observed in comparative analysis of *B. bassiana* strains interacting with the domestic silk moth *Bombyx mori* epicuticle (Wang *et al*., 2017). However, our analysis revealed that ILB308 instead up-regulated genes associated with RNA biosynthesis, DNA repair, and genetic information processing. A similar trend was reported when comparing ILB308 with the hypovirulent strain ILB299 (Sessa *et al*., 2024) whereas previous studies suggest that reduced virulence in *B. bassiana* is linked to down-regulation of DNA replication and energy metabolism (Jirakkakul *et al*., 2018). This implies that ILB308’s increased virulence may be linked to a more efficient transcriptional response and cellular proliferation rather than specific hydrocarbon degradation pathways. While this association is intriguing, this results suggest a range of general processes that may be pleiotropic and, consequently, do not directly imply specific virulence factors to be linked to virulence difference to be used for enhancement of low-virulence strains through genetic engineering.

To refine our selection of candidate genes for virulence enhancement, we combined transcriptomic data from epicuticle interaction with previously identified genes induced during infection at 4 dpi. This approach allowed us to pinpoint genes that are not only responsive to epicuticle contact but also relevant in later infection stages. Among these, we identified *cpf* and *Blys8*, two well-characterized virulence factors previously associated with immune evasion in the hemocoel (Cen *et al*., 2017; Mou *et al*., 2022). Interestingly, our data shows that these genes are also up-regulated in ILB308 during epicuticle interaction, suggesting a broader role in fungal pathogenicity and a possible key difference in virulence between ILB308 and ILB205. Also, we wanted to highlight ILB308’s enhancing of virulence through HC15 pre-growth. Based on this, we further refined our selection by identifying genes that were both exclusively up-regulated in ILB308 in HC15_MM+EC condition (and not in MM or MM+EC) and predicted to be secreted, as secreted proteins often function as virulence factors in entomopathogens (Wang and Wang, 2017; Griffiths *et al*., 2018; Mou *et al*., 2021, 2022). This selection led to the identification of five secreted candidate genes. Among them, *BbCBM9_1* was the only one containing a carbohydrate-binding module (CBM9), a domain well-documented in bacterial systems for its role in polysaccharide recognition and hydrolysis (Guillén *et al*., 2010). Although CBM9-containing proteins have not been extensively studied in fungal pathogens, their role in recognizing and hydrolyzing polysaccharides like cellulose, hemicellulose, and xylan is well-established in bacteria such as *Thermotoga maritime* (Guillén *et al*., 2010). Interestingly, a similar secreted CBM9 domain-containing protein (*CPF28*) expression was analyzed by western-blot analysis in the pathogenic fungus *C. posadasii* and found to be up-regulated in response to N-acetylglucosamine (GlcNAc) (Lunetta *et al*., 2010; Lunetta and Pappagianis, 2014), the monomer of chitin and a product of insect cuticle degradation (Zhang *et al*., 2024). Additionally, a homolog of *BbCBM9_1* in *B. bassiana* ARSEF2860 was verified as secreted using a secretome analysis and up-regulated after exposure to cattle tick *Rhipicephalus microplus* cuticle (Santi *et al*., 2018). These findings suggest that *BbCBM9_1* is secreted and involved in epicuticle interaction, potentially recognizing chitin or its derivatives, reinforcing its role as a candidate for enhancing virulence in *B. bassiana*.

Later, we overexpressed *BbCBM9_1* in ILB205 to assess whether its overexpression enhances virulence, one of the main objectives of this study. Our results showed that the overexpression mutants exhibited a significant increase in virulence against *P. guildinii* and *T. molitor*, along with improvements in germination, growth on different axenic media, and tolerance to oxidative and cell wall stressors. Since the *CPF28* ortholog in *C. posadasii* has been linked to increased chitinolytic activity upon induction (Lunetta *et al*., 2010; Lunetta and Pappagianis, 2014), *BbCBM9_1* may facilitate more active/efficient cuticle degradation, and enhancing the release of GlcNAc that later can become an integral part of the fungal cell wall, increasing tolerance to for example cell wall stresses as reported previously in *B. bassiana* (Zhang *et al*., 2024). We suggest that virulence improvement could lay in improved cell wall and oxidative stress resistance, augmented host defense evasion and more efficient cuticular degradation.

In conclusion, our study demonstrates that *BbCBM9_1* plays a key role in enhancing *B. bassiana* virulence and holds promise for improving fungal strains used in biocontrol of *P. guildinii*. Our omics approach revealed that ILB308 activates immune evasion and specialized metabolic responses during infection, and that differences in BbCBM9_1 expression between strains may be driven by gene interactions or epigenetic modifications. From a technological point of view, these findings validate the use of integrated transcriptomic and genomic methods to identify novel virulence genes and enhance pest management strategies.

## Data availability statement

RNA-seq datasets generated during the current study are available at NCBI BioProject repository, under accession number PRJNA1117794.

## Acknowledgements

This project was supported by the National Institute of Agricultural Research of Uruguay (INIA) project SA-24, and by project FMV_1_2021_1_169591 from the Agencia Nacional de Investigación e Innovación (ANII) of Uruguay. HO Scholarships were founded by INIA, ANII, Programa de Desarrollo de las Ciencias Basicas (PEDECIBA) of Universidad de la República, Uruguay and Comisión Sectorial de Investigación Científica (CSIC) of Uruguay. The funding bodies played no role in the design of the study and collection, analysis, and interpretation of data and in writing the manuscript. Authors acknowledge the use of ChatGPT (https://chatgpt.com/) for assistance in grammar and language correction.

---

**Supplementary Figure 1.**
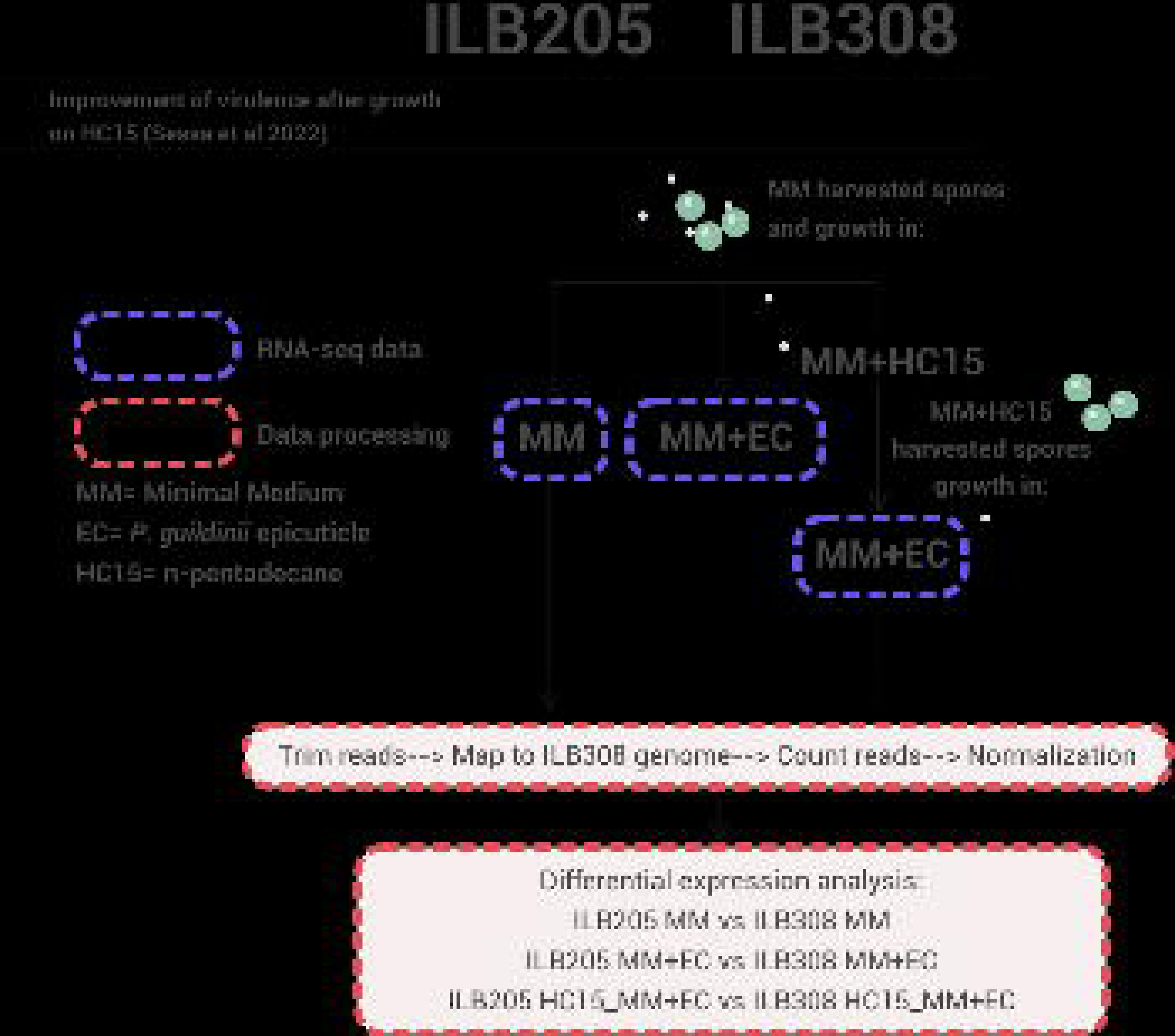
Data collection and analysis workflow. Diagram displaying the sample collection of three different axenic conditions analyzed in this study and how the expression analysis was conducted. MM: Minimal medium; MM+EC: MM supplemented with P. guildinii epicuticle; HC15_MM+EC: conidia pre grown in MM supplemented with n-pentadecane (HC15), harvested and then grown in MM+EC.

**Supplementary Figure 2.**
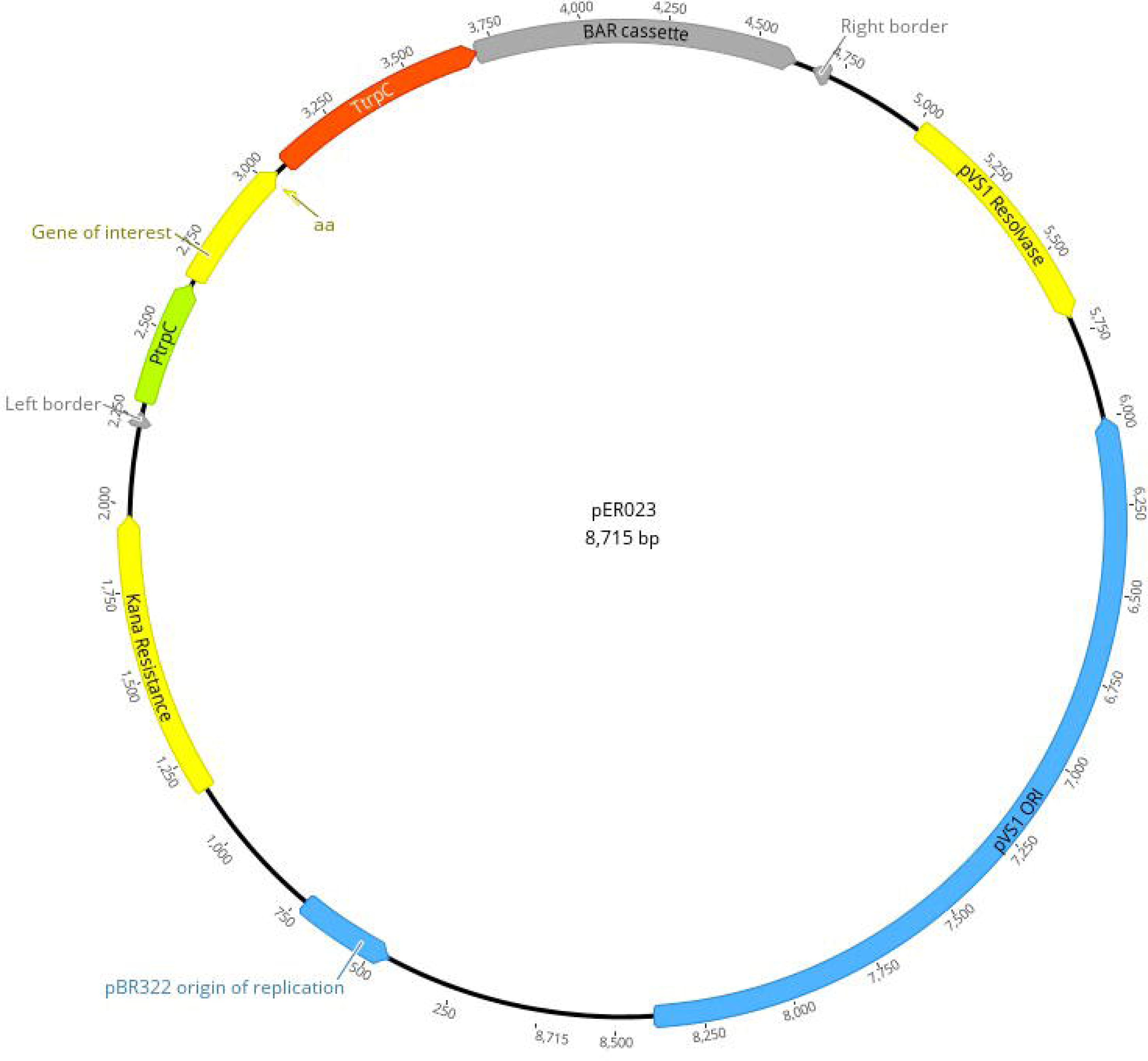
Plasmid generated in this work. Schematic representation of the binary vector pERO23 containing a gene of interest inserted at the EcoRV site, designed for fungal overexpression using *Agrobacterium tumefaciens*-mediated transformation. The vector includes a bar cassette conferring resistance to phosphinothricin and a kanamycin (Kana) resistance gene for bacterial selection. Expression is driven by the *Aspergillus nidulans* TrpC promoter (PTrpC) and terminated by the TrpC terminator (TtrpC). The Right and left border indicate the T-DNA limits required for transfer into the fungal genome

**Supplementary Figure 3.**
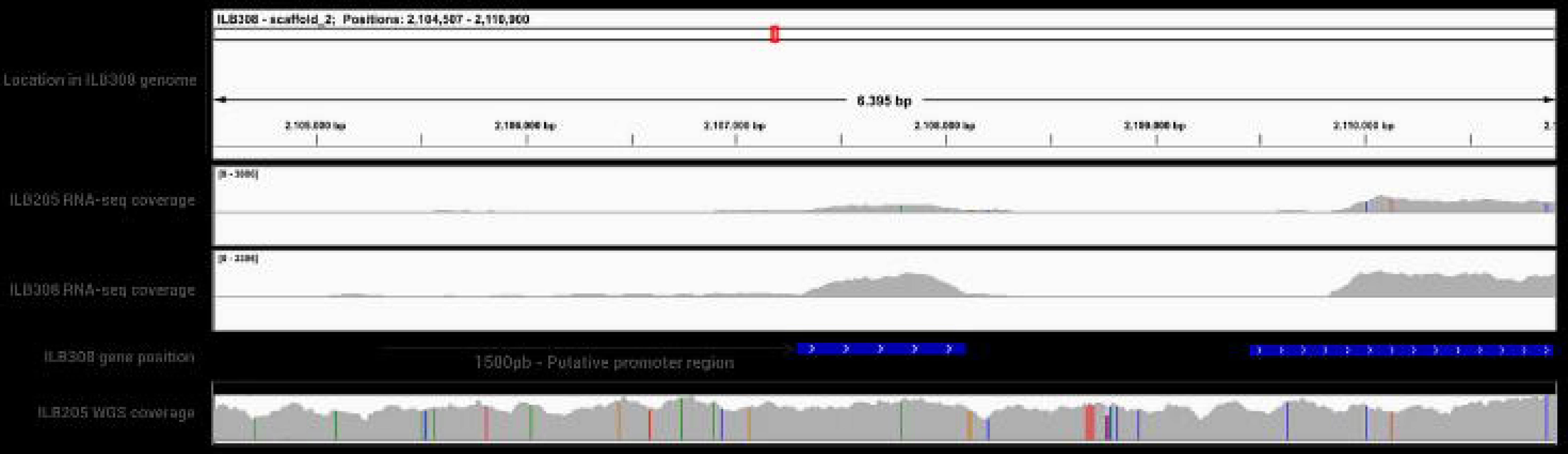
DNA and transcriptome mapping reveal sequence differences between ILB308 and ILB205 around BbCBM9_1. From top to bottom, the five tracks represent: i) The genomic location of the region in the ILB308 genome, ii) Coverage plot of an example of RNA-seq from ILB205 under HC15_MM+EC conditions; iii) Coverage plot of an example of RNA-seq from ILB308 under HC15_MM+EC conditions, iv) Gene annotation for the region, displaying BbCBM9_1, the adjacent downstream gene, and the putative promoter Region of BbCBM9_1; v) DNA-seq coverage from ILB205 mapped to ILB308 genome, SNPs between ILB205 and ILB308 are highlighted with different colors (red: T; green: A; blue: C; orange: G).

**Supplementary Figure 4.**
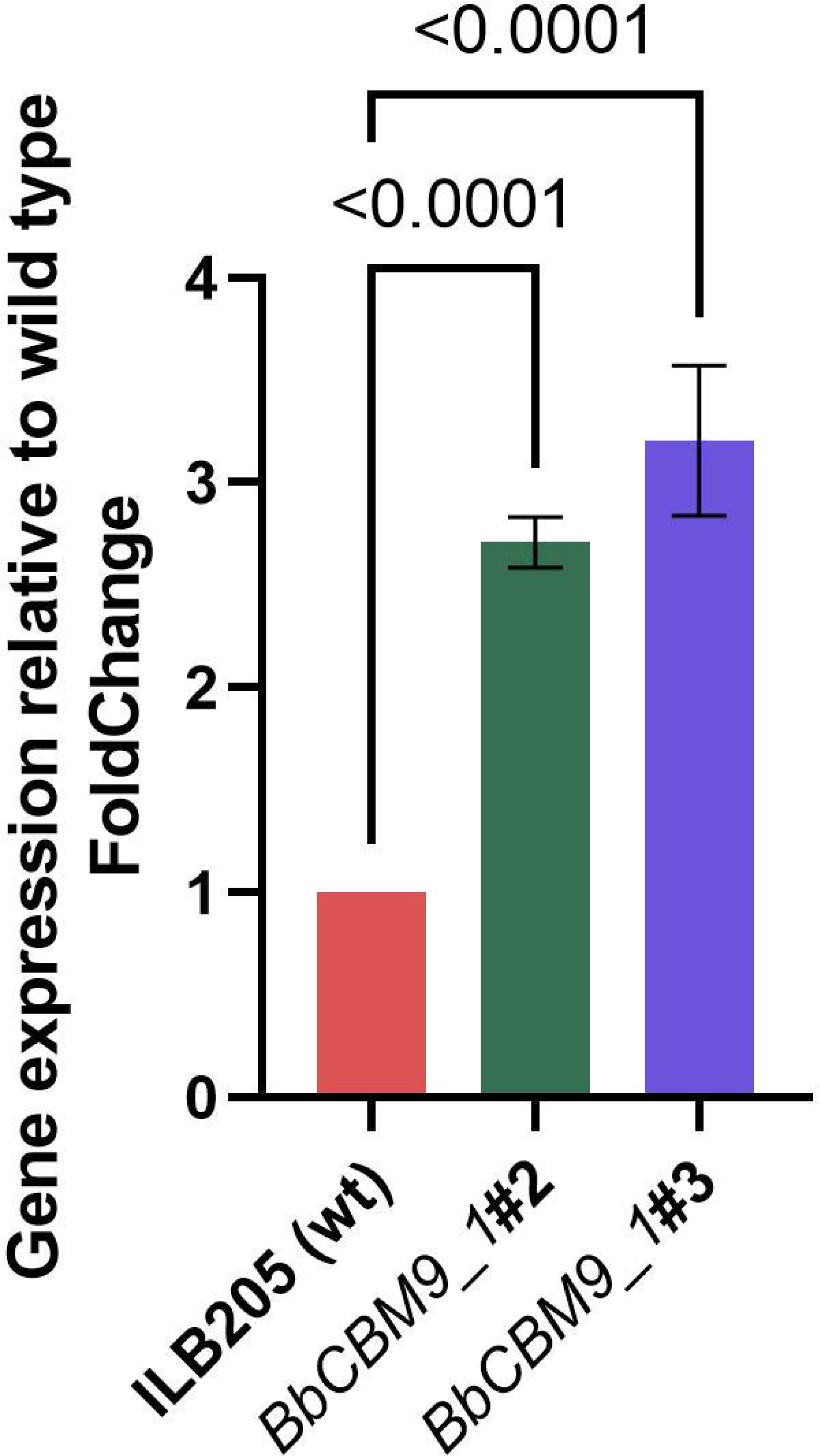
Relative expression levels of *BbCBM9_1* in overexpression mutants compared to the wild-type strain ILB205 under axenic conditions (PDA). GAPDH was used as a reference gene. n_wildtype_=3, n_BbCBM9_1#2_=3, n_BbCBM9_1#3_=3. *P-values* < 0.01 according to a one-way ANOVA followed by a Tukey posthoc test of results between overexpression lines versus the wild type are displayed on the plot based on triplicate experiments. Error bars: standard errors (SEs) of the means from three independent replicates.

